# Co-evolution of gene transfer agents and their alphaproteobacterial hosts

**DOI:** 10.1101/2023.08.11.553018

**Authors:** Roman Kogay, Olga Zhaxybayeva

## Abstract

Gene transfer agents (GTAs) are enigmatic elements that resemble small viruses and are known to be produced during nutritional stress by some bacteria and archaea. The production of GTAs is regulated by quorum sensing, under which a small fraction of the population acts as GTA producers, while the rest become GTA recipients. In contrast to canonical viruses, GTAs cannot propagate themselves because they package pieces of the producing cell’s genome. In alphaproteobacteria, GTAs are mostly vertically inherited and reside in their hosts’ genomes for hundreds of millions of years. While GTAs’ ability to transfer genetic material within a population and their long-term preservation suggests an increased fitness of GTA-producing microbes, the associated benefits and type of selection that maintains GTAs are poorly understood. By comparing rates of evolutionary change in GTA genes to the rates in gene families abundantly present across 293 alphaproteobacterial genomes, we detected 59 gene families that likely co-evolve with GTA genes. These gene families are predominantly involved in stress response, DNA repair, and biofilm formation. We hypothesize that biofilm formation enables the physical proximity of GTA-producing cells, limiting GTA-derived benefits only to a group of closely related cells. We further conjecture that population structure of biofilm-forming sub-populations ensures that the trait of GTA production is maintained despite the inevitable rise of “cheating” genotypes. Because release of GTA particles kills the producing cell, maintenance of GTAs is an exciting example of social evolution in a microbial population.

**Importance:** Gene transfer agents (GTAs) are viruses domesticated by some archaea and bacteria as vehicles for carrying pieces of the host genome. Produced under certain environmental conditions, GTA particles can deliver DNA to neighboring, closely related cells. Function of GTAs remains uncertain. While making GTAs is suicidal for a cell, GTA-encoding genes are widespread in genomes of alphaproteobacteria. Such GTA persistence implies functional benefits but raises question about how selection maintains this lethal trait. By showing that GTA genes co-evolve with genes involved in stress response, DNA repair, and biofilm formation, we provide support for the hypothesis that GTAs facilitate DNA exchange during the stress conditions and present a model for how GTAs persist in biofilm-forming bacterial populations despite being lethal.

## Introduction

Multiple bacteria and archaea produce <u>G</u>ene <u>T</u>ransfer <u>A</u>gents (GTAs) – the viriforms whose function and mode of selection to maintain them remain unsolved (1–3). These domesticated virus-derived elements are encoded by genes in their host’s genome and, when produced, resemble tailed double-stranded DNA (dsDNA) viruses (phages). In contrast to viruses, GTAs do not package the genes that encode them, and instead contain fragments of the producing host’s genome (1, 4–7). Experimentally, GTAs are most studied and characterized in the alphaproteobacteria *Rhodobacter capsulatus* (RcGTA) and *Caulobacter crescentus* (6, 8, 9), but they are also produced by several additional bacterial and archaeal species (10). Many more prokaryotes encode GTA-like genes (5, 11–18), and the presence of GTA-like genes in almost 60% of publicly available alphaproteobacterial genomes (19) suggests that GTA production is more widespread than currently appreciated.

RcGTA production is a population-level phenomenon: it is triggered by nutrient depletion (20) and is regulated by quorum sensing (21). Only a small subset of the population acts as RcGTA producers; the remaining cells become recipients, by displaying specific polysaccharide receptors for RcGTAs adsorption (22) and expressing competence genes (23). Genetic pieces delivered by RcGTAs to a recipient cell can be integrated into the cell’s genome via homologous recombination (24).

The benefits of GTA production and of acquiring GTA-packaged DNA in a microbial population are not fully understood. Since their discovery, GTAs were hypothesized to mediate DNA repair (25), and recently this hypothesis was confirmed by experimental demonstration of GTA-mediated DNA repair via homologous recombination in *C. crescentus* (6). Moreover, facilitation of DNA damage repair appears to improve the survival of *C. crescentus* populations in nutrient limited conditions (6), possibly due to a reduction in mutational load. Beyond the repair of already existing genes, released GTA particles could enable exchange of beneficial traits in a microbial population (8, 26) and provide nutrients to surrounding cells as the programmed cell death phenomenon does (27, 28), although these hypotheses remain to be experimentally verified. Despite these putative population-level benefits, GTA-producing cells lyse and therefore leave no progeny, making it impossible for selection that maintains GTA production to act on the level of individual cells. Better understanding of GTA production and reception cycle and of genes underlying it will likely help us elucidate ecological role of GTAs in microbial communities, and details of the population-level selection that preserves the trait.

The RcGTA is encoded and regulated by at least 24 genes that are distributed across five different loci (29). Seventeen genes are located in one locus that is commonly referred as the head-tail cluster (2) (**Figure 1A**). The locus encodes the majority of structural proteins required for the RcGTA particle assembly (30). Products of many additional “host” genes are critical for the regulation of the RcGTA particle production, DNA uptake, and DNA integration. For example, the CckA-ChpT-CtrA phosphorelay system, which controls the cell cycle and DNA replication in other model organisms (31–33), modulates production of RcGTA particles and their release (34, 35). Serine acetyltransferase (cysE1), which is required for biofilm formation, plays a critical role for the optimal receipt of RcGTAs (36). Capsular polysaccharides, which serve as RcGTA receptors, are synthesized under control of GtaR/I quorum-sensing genes (22). Competence machinery proteins ComEC, ComF and ComM facilitate entry of DNA into cells (23). Integration of the incoming genetic material into the host genome via homologous recombination is facilitated by DprA and RecA (24). It is likely that products of multiple other “host” gene families important for the proper functioning of GTAs remain to be discovered, and in this study we use a comparative genomics approach to search for such genes.

**Figure 1.**
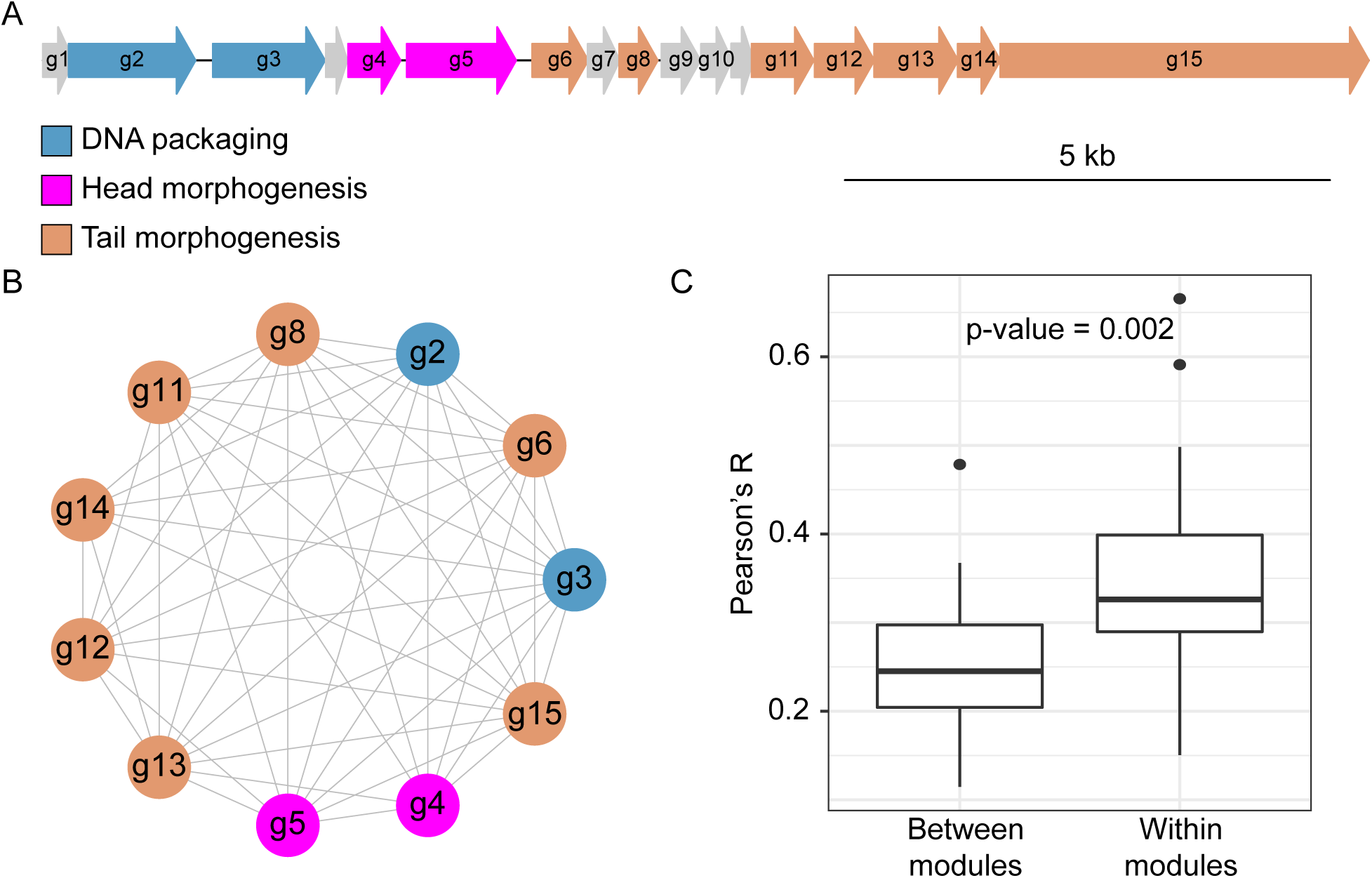
Co-evolution of 11 reference GTA genes from head-tail clusters. (**A**) The head-tail cluster of the RcGTA, as encoded in *R. capsulatus* strain SB 1003 genome (GenBank accession CP001312.1; spanning locus tags RCAP_rcc01682 – RCAP_rcc01698). Each gene is represented by an arrow and is drawn proportional to its length (see scale bar). Genes shown in non-gray color correspond to the 11 GTA reference genes used in the ERC analysis. Reference genes from each functional module are depicted in a distinct color. **(B)** Co-evolution network of the reference GTA genes. Nodes represent genes and are connected by edges if there is a significant evolutionary rate covariation (Pearson’s R > 0 and p-value <0.05 after Bonferroni correction). **(C)** Comparison of the rate covariation strength in reference GTA genes that belong either to different functional modules (“Between Modules”) or to the same functional module (“Within Modules). Boxplots represent median values that are bounded by first and third quartiles. Whiskers illustrate data points that lie within 1.5 * interquartile range. Significance was measured using Mann-Whitney U-test.

Genes that are involved in the similar molecular processes, or co-expressed together, tend to co-evolve with each other (37, 38), and, vice versa, the protein-protein interactions can be unveiled by finding co-evolving genes that encode the interacting proteins (39, 40). The co-evolution among genes can be effectively identified via Evolutionary Rate Covariation (ERC) approach (37, 41, 42). Evolutionary rate covariation measures the degree of correlation of changes in evolutionary rates across the phylogenies of a pair of proteins, assuming that functionally related proteins have similar selection pressures, resulting in coordinated changes in substitution rates (37, 38). Because GTA head-tail cluster has resided within GTA-containing alphaproteobacterial genomes for at least 700 million years (13) and, as mentioned above, GTA production is tightly integrated into the molecular circuits of the GTA-carrying bacteria, evolutionary rates of gene families involved in GTA lifecycle are expected to correlate with the rates of the GTA genes.

Head-tail cluster genes are easily detectable across genomes in a large clade of alphaproteobacteria (11, 13) and therefore provide a rich dataset for comparative analyses of evolutionary rates. In this study, we examined the evolutionary rate covariation patterns of protein-coding genes encoded in 293 representative alphaproteobacterial genomes that contain either complete or nearly complete GTA head-tail clusters. We found that GTA head-tail cluster genes co-evolve with 59 gene families, 55 of which have not been previously linked to GTAs. Thus, we dramatically expand the list of genes that could be important for GTAs’ functionality and hence could provide insights into GTAs’ role in bacterial populations. By combining our findings with the existing knowledge about GTAs, we propose a model that explains persistence of GTA production in bacterial populations.

## Results

### Alphaproteobacterial GTA genes co-evolve with each other

The ERC method was developed for and tested on eukaryotic genes (37, 38) and, to our knowledge, has not been applied to bacterial genomes. Therefore, before using the approach to identify genes co-evolving with GTA genes, we evaluated it on GTA genes found in the 293 alphaproteobacterial genomes. Because genes in the GTA head-tail cluster have a common promoter (8, 43) and the gene products are functionally related (i.e., produce a GTA particle that has DNA packaged into its head), we expected strong co-evolution among GTA genes. Indeed, we found that 51 out of 55 possible pairs of the 11 reference GTA genes (see **Methods** for definition) have significantly similar co-variation of evolutionary rates, as measured by the Pearson’s coefficient (p-value < 0.05 after Bonferroni correction). Each reference GTA gene co-evolves with at least 7 other reference GTA genes, with 6 of them co-evolving with all 10 other reference GTA genes (**Figure 1B**). These findings suggest that the ERC method adequately identifies co-evolving genes in alphaproteobacterial genomes.

### Co-evolving alphaproteobacterial genes encode functionally related proteins

Co-evolving genes of eukaryotes identified through ERC analyses were shown to be either functionally related or involved in similar biological processes (37, 38). To examine how robustly the ERC method can identify functionally related gene pairs in our dataset of alphaproteobacterial genomes, we (i) evaluated rates of covariation within functional modules of the GTA head-tail cluster and (ii) examined a relationship between co-evolution and literature– and experiment-based functional inferences for a subset of gene families nearly universally found across GTA-containing alphaproteobacteria.

GTA head-tail cluster encodes three modules that are responsible for distinct functional stages of GTA production: DNA packaging, head morphogenesis, and tail morphogenesis (2, 29, 30) (**Figure 1A**). Phage genes within the same functional class are more likely to interact with one another (44). We found that the Pearson’s correlation coefficient is significantly higher (and, therefore, co-evolutionary signal is significantly stronger) for reference GTA genes within each module than between the reference GTA genes from different modules (Mann-Whitney U-test, p-value = 0.002) (**Figure 1C**). These findings suggest that the strength of co-evolutionary signal measured by the ERC analysis correlates with the degree of physical and functional interactions among GTA genes.

Expanding our analysis beyond GTA genes, we examined protein-coding genes in a model marine bacterium *Phaeobacter inhibens*, whose genome encodes the largest number (1,370) of genes from 1,470 gene families nearly universally found across GTA-containing alphaproteobacteria, and in *Caulobacter crescentus* and *Dinoroseobacter shibae*, which are experimentally shown to produce GTAs (5, 6). The information about the interactions of most proteins encoded by the protein-coding genes in the genomes of these bacteria is available in the STRING database (45), and the experimental relative fitnesses of many genes are cataloged in Fitness Browser (46). Using the ERC analysis on the 1,320, 1,199, and 1,255 genes, we identified 10,514, 9,373 and 9,675 co-evolving gene pairs for *P. inhibens*, *C. crescentus*, and *D. shibae*, respectively (**Figure S1**). The co-evolution networks, in which nodes correspond to genes and edges designate the presence of significant evolutionary rate covariation (see FigShare repository), are significantly similar to both the networks of the pairwise interactions of the encoded proteins and the networks of fitness effects for all three genomes (p-value < 0.001, permutation test) (**Figure 2**). Moreover, co-evolving genes are more likely to belong to the same Clusters of Orthologous Groups (COG) category (assortativity of 0.096 for *P. inhibens*, 0.083 for *C. crescentus*, and 0.094 for *D. shibae*; p-value < 0.001 in permutation tests for all three genomes) (**Figure 2**). These comparisons show that co-evolving genes that encode well-characterized proteins (defined as present in the STRING and COG databases) and a subset of genes needed for specific environmental conditions (as determined by the Fitness Browser database) indeed tend to encode functionally related proteins. For example, *P. inhibens’* gene pairs with the two largest Pearson’s coefficients encode proteins that are involved in the same biological processes (**Figure 3**): *imuB* and *dnaE2* genes (Pearson’s coefficient r = 0.800) are located in the same operon and are involved in SOS-induced mutagenesis and translesion synthesis (47, 48), while *addA* and *addB* genes (Pearson’s r = 0.735) are known to assemble into the heterodimeric complex to facilitate homologous recombination and DNA repair (49, 50).

**Figure 2.**
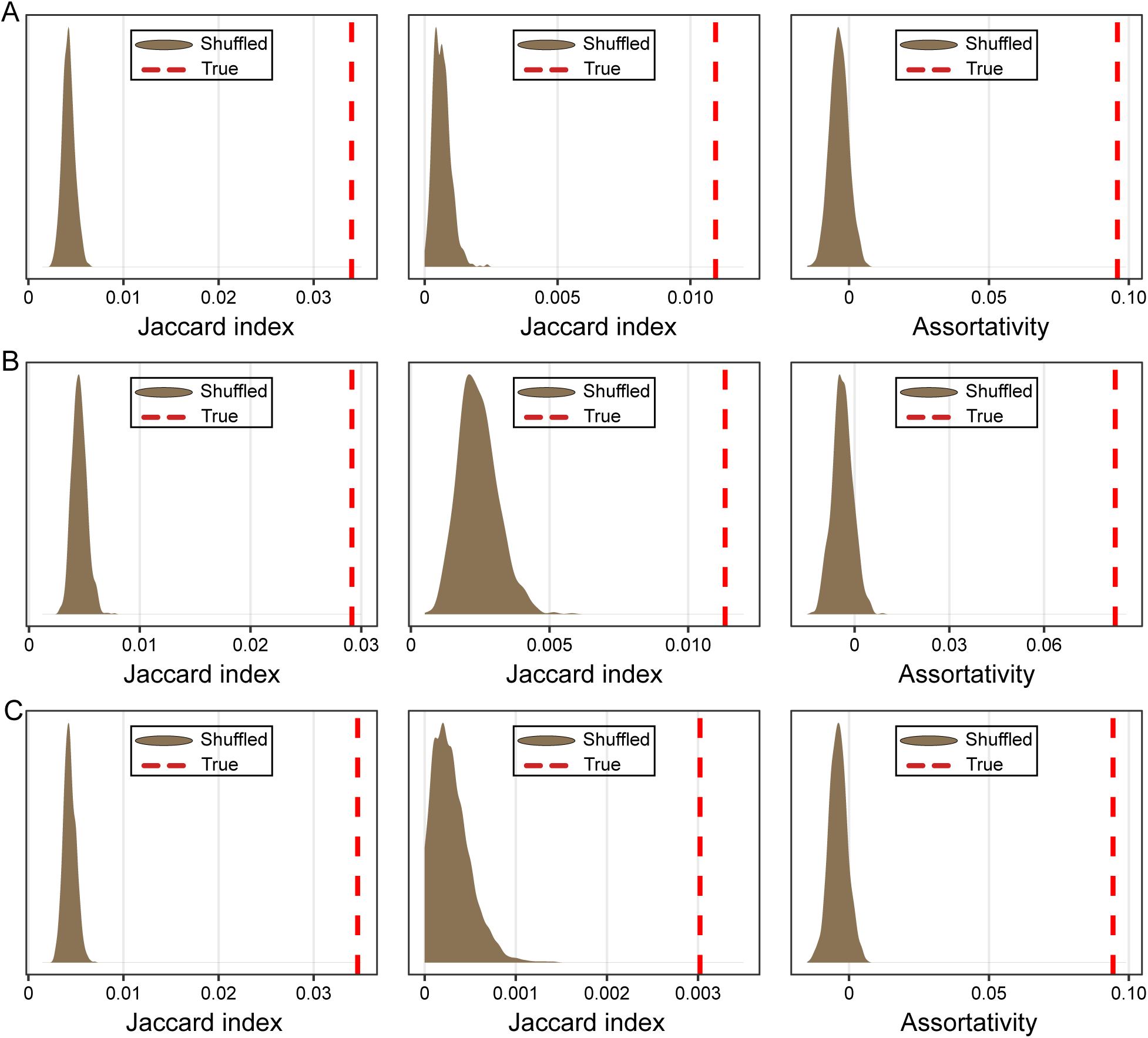
Strengths of correlations between the covariation evolutionary rate of genes and function of the proteins the genes encode, as measured by permutation tests. The analyses were carried out for (A) *Phaeobacter inhibens*, (B) *Caulobacter crescentus*, and (C) *Dinoroseobacter shibae*. Graphs on the left show the comparisons between the co-evolution and PPI networks, while graphs in the middle show the comparisons between co-evolution and co-fitness network. For these graphs, the distributions in brown represent distances between the PPI/co-fitness networks and 1,000 co-evolution networks, in which edges were randomly shuffled. The dashed red lines indicate the Jaccard index from the non-shuffled network comparisons. Graphs on the right show positive assortativity between co-evolution and COG functional category assignment. The distributions in brown represent assortativity coefficient values for 1,000 networks, in which COG labels were randomly shuffled. The dashed red lines indicate the assortativity coefficient of the non-shuffled network.

**Figure 3.**
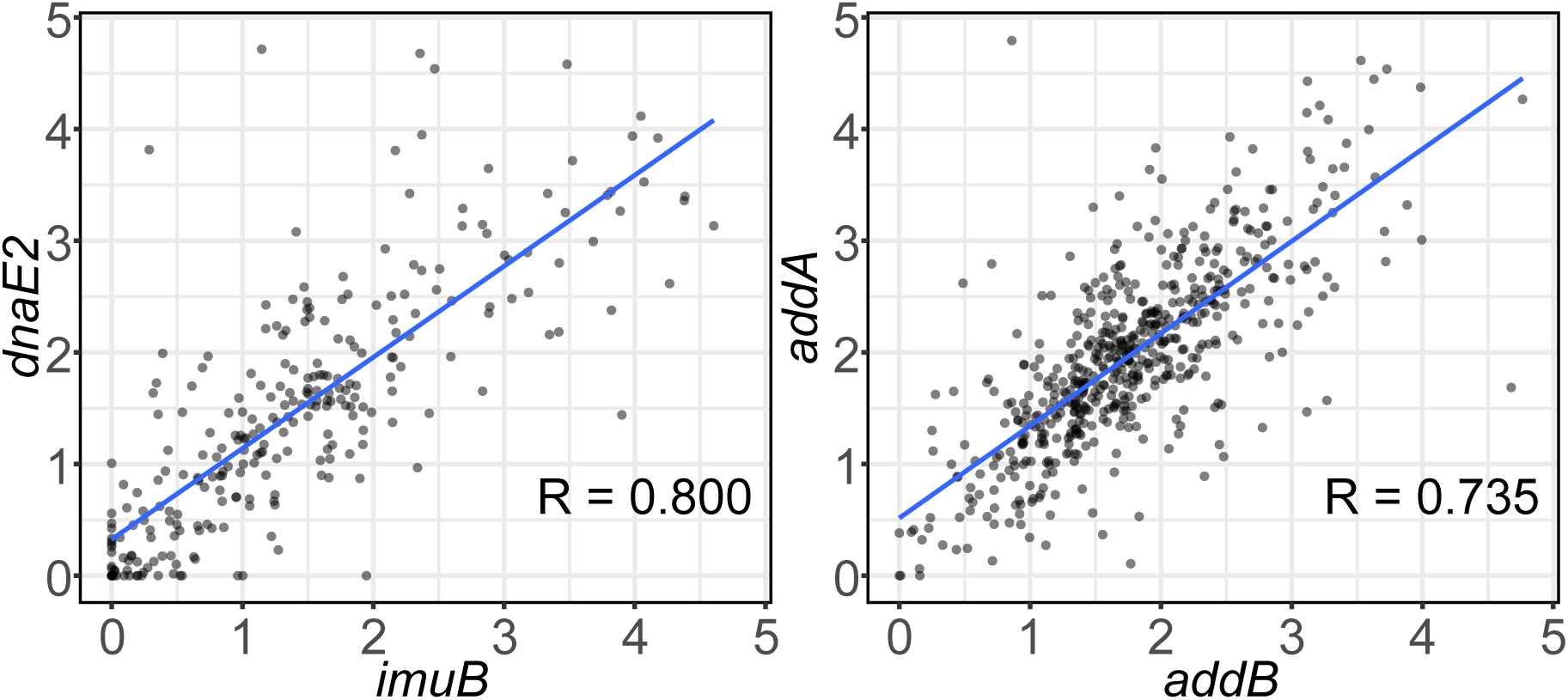
Covariation of evolutionary rates for two gene pairs in *P. inhibens* with the largest Pearson’s coefficients. Each dot represents the normalized branch length of the proteins shown on x– and y-axes. Blue line depicts the linear regression line. The value of Pearson’s R is shown on each panel.

It is worth noting that some of the genes identified as co-evolving in our analysis are not designated as encoding interacting proteins in the STRING database. However, given incompleteness of our knowledge about functionality of proteins encoded in a bacterial genome, these genes may represent functional connections yet unidentified in STRING. Indeed, ERC analysis has been used to uncover novel protein-protein interactions, especially between hypothetical proteins (40, 51, 52). Here are two examples of proteins identified as co-evolving in our analyses and likely interacting based on what’s known about their functions, but not designated as such in the STRING database. The *yfgC* gene, which encodes a periplasmic metalloprotease involved in assembly of outer membrane proteins (53), co-evolves with both the *lptD* and *bamB* genes (**Figure 4**). The YfgC protein plays a crucial role in the assembly of LptD, an outer membrane protein that participates in the lipopolysaccharide assembly (53). The YfgC also interacts with the β-barrel-assembly machinery (BAM) complex, which consists of four lipoproteins, including BamB, and facilitates the assembly and integration of proteins into the outer membrane (53, 54).

**Figure 4.**
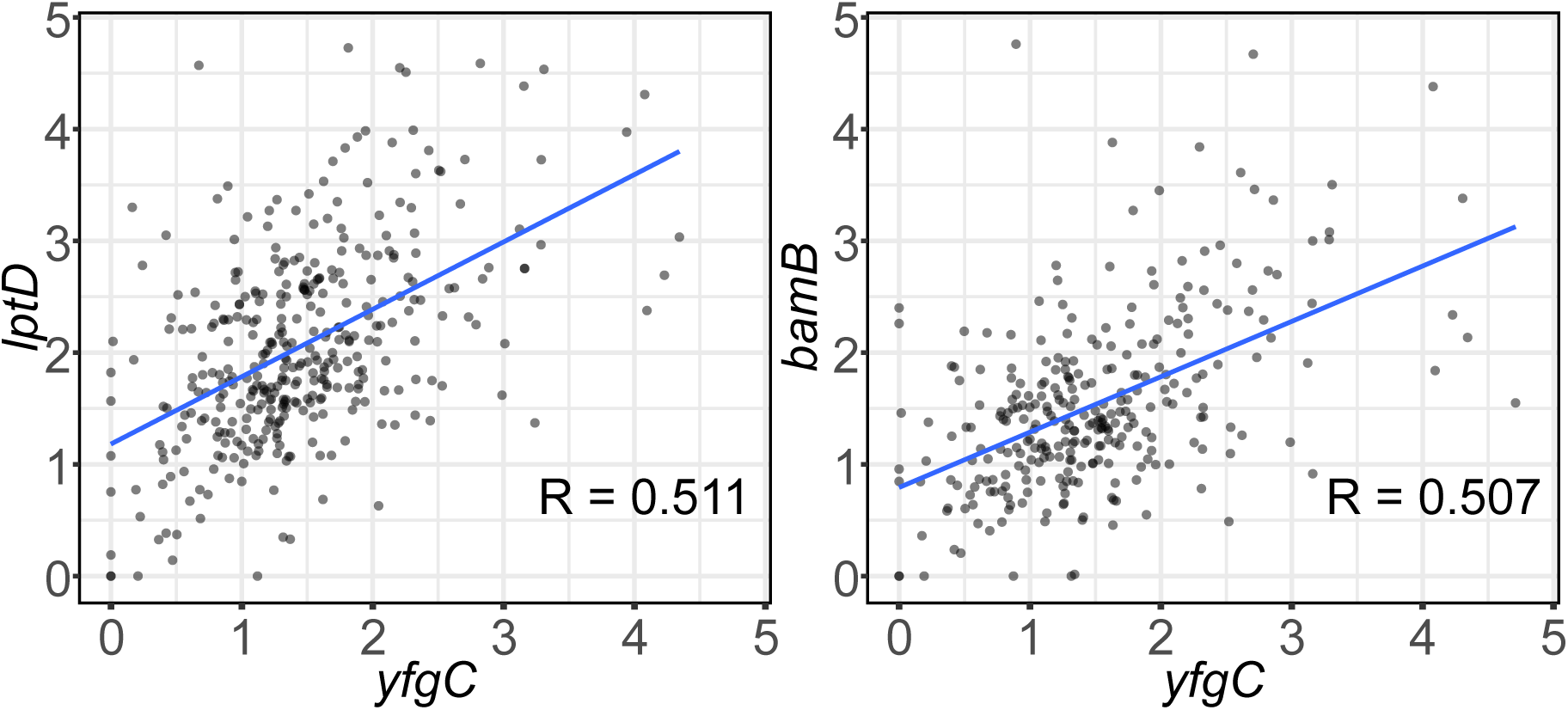
Covariation of evolutionary rates for two gene pairs in *P. inhibens* with no evidence of interactions for their protein products in the STRING database. Each dot represents the normalized branch length of the proteins shown on x– and y-axes. Blue line depicts the linear regression line. The value of Pearson’s R is shown on each panel.

### GTA genes co-evolve with at least fifty-nine other gene families

By analyzing 1,470 gene families almost universally present among the 293 representative GTA-containing alphaproteobacteria, we identified 59 gene families that co-evolve with at least 5 reference GTA genes (**Table S1**) (see **Methods** for selection criteria).

Notably, four of the 59 gene families – encoding tail fiber protein (DUF2793), competence proteins (comEC and comF), and DNA-protecting protein that facilitates homologous recombination (dprA) (**Table 1**) – have already been shown to play important roles in the RcGTA lifecycle (23, 24, 29). Because tail fiber proteins are a part of the RcGTA particle and thus physically interact with other RcGTA proteins (29), DUF2793’s co-evolution with other structural GTA genes is expected. The protein products of the remaining three genes (ComEC, ComF, and DprA) are required for the acquisition of DNA delivered by RcGTAs, and interact physically only with DNA molecules (23, 24). This finding demonstrates that the ERC approach indeed could predict genes functionally linked to the GTA lifecycle.

**Table 1.**
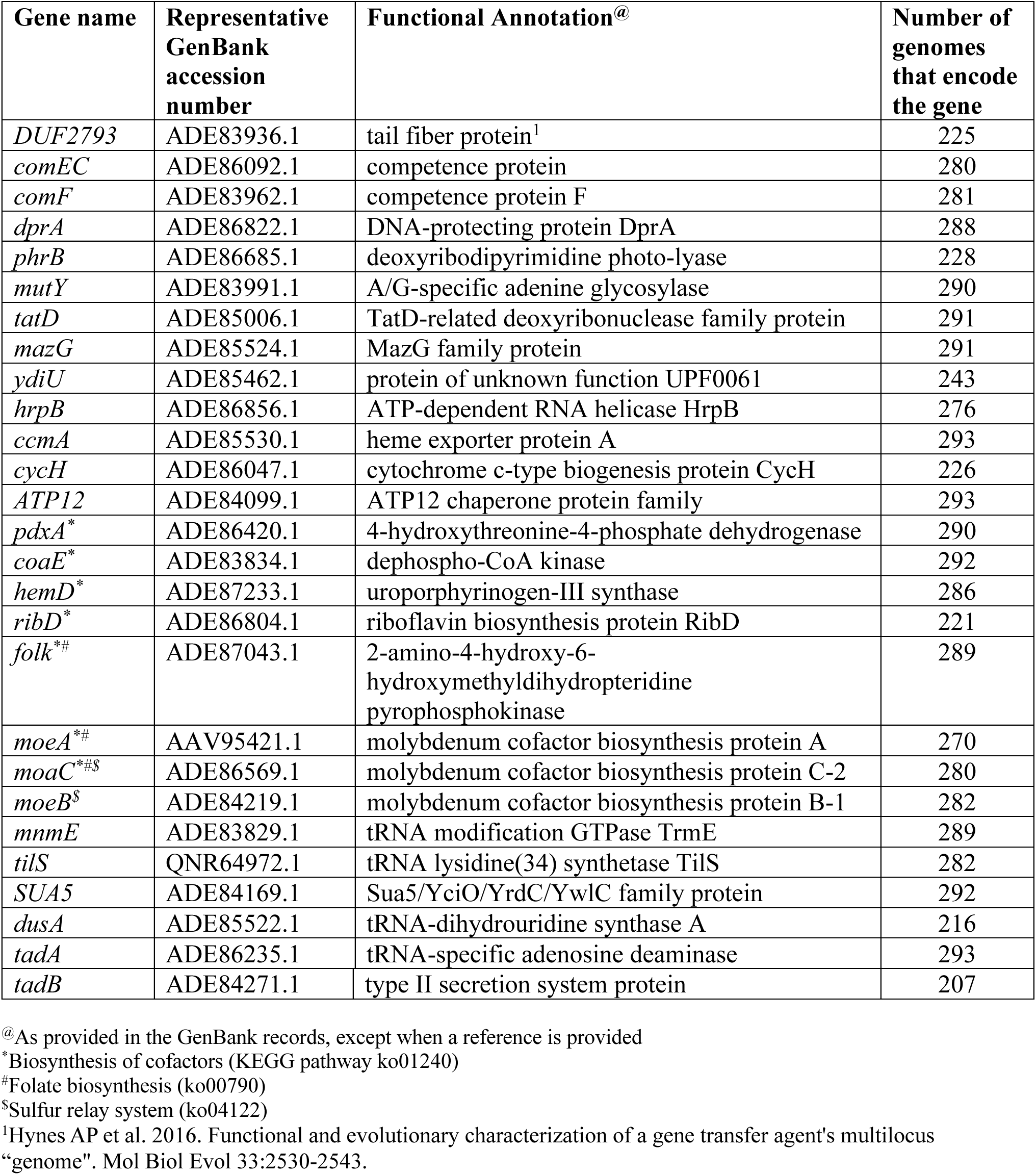
Twenty-seven gene families that co-evolve with reference GTA genes and discussed throughout the manuscript.

The remaining 55 gene families are involved in various functions (**Tables 1, 2** and **S1**). While 32 of the 55 gene families (**Table 2**) offer exciting opportunities for future research into GTA lifecycle, 23 gene families (**Table 1**) can be either directly linked to the GTA lifecycle by being involved in DNA repair or are likely to be under similar selection pressure as GTA genes due to shared ecological importance (stress response, biofilm formation, oxidative respiration, and cofactor biosynthesis), as elaborated below.

**Table 2.**
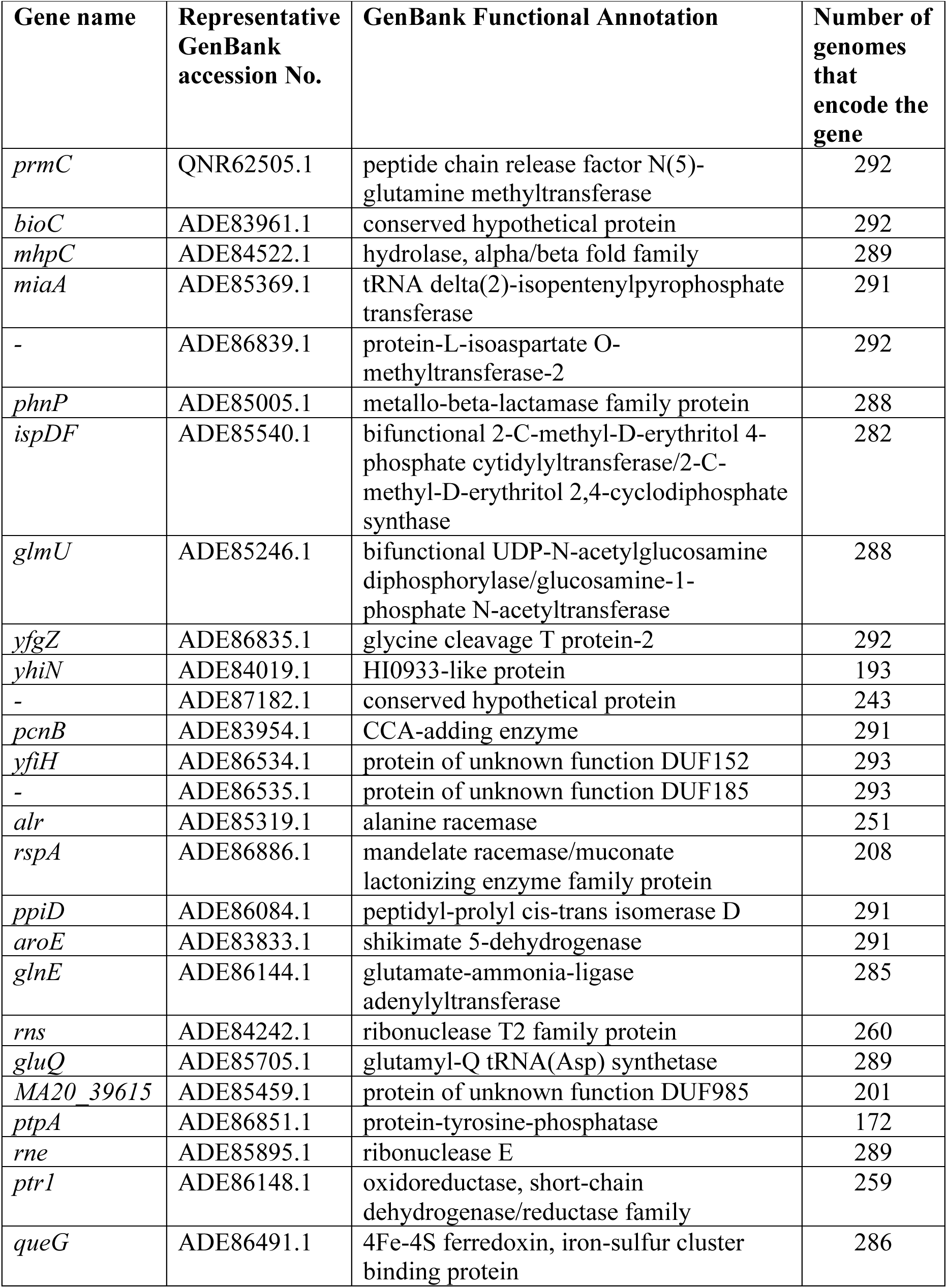
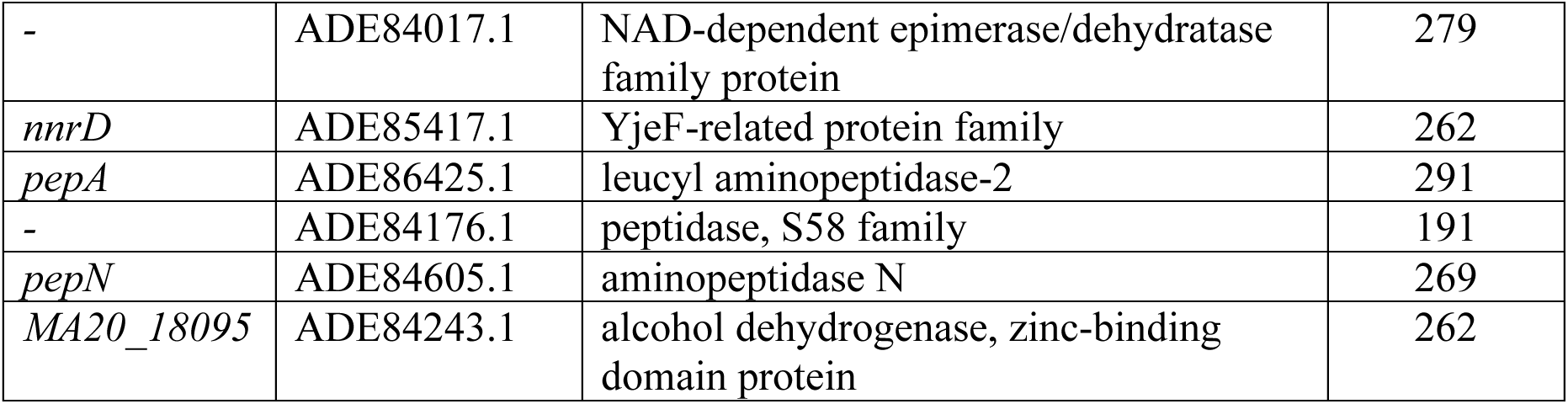
Thirty-two gene families that co-evolve with reference GTA genes, but without known connection to the GTA production cycle.

Three of the 22 genes – *mutY* (encoding a glycosylase), *phrB* (encoding a DNA photolyase), and *tatD* (encoding an exonuclease) – play roles in repair of DNA damage induced by various oxidative agents and UV light (55–57). Glycosylase actively modulates homologous recombination (58), which could be important for facilitating integration of GTA-derived genetic material into the recipient’s genome (6, 9). Photolyase and tatD exonuclease do not directly participate in homologous recombination, but are involved in DNA repair process (6, 55, 56) and thus are likely to be under the similar selection pressure as GTA genes.

Two genes (*mazG* and *ydiU*) identified in our screen are involved in stress response. The product of the *mazG* gene modulates the programmed cell death in *Escherichia coli* and regulates intracellular level of ppGpp, the universal ‘alarmone’, which was previously implicated in the regulation of GTAs production (20, 59). The ppGpp molecule is involved in a cellular response to a variety of stress conditions, including nutritional stress. The product of the *ydiU* gene mediates UMPylation of bacterial chaperones, improving bacterial fitness under the stressful environmental settings (60).

Thirteen genes identified in our analyses have relevance to biofilms. While these genes are linked to the formation of biofilms, they are also involved in a range of other molecular functions. The *hrpB* gene, which encodes ATP-dependent RNA helicase, has been shown to be important for both biofilm formation and adhesion on surfaces (61). The *ccmA* gene, *cycH* gene, and a gene from COG5387 family (*ATP12*) are involved in oxidative respiration, which promotes bacterial survival in the biofilms (62, 63). Moreover, as a group, 59 co-evolving gene families are enriched in three metabolic pathways relevant to biofilms: cofactor biosynthesis, folate biosynthesis and sulfur relay system (hypergeometric test, p-value < 0.05, Bonferroni correction) (**Table 1**). Two genes from the ‘cofactor biosynthesis’ pathway (*moeA* and *moaC*) are involved in the molybdenum cofactor biosynthesis. Both metabolism of folate and molybdenum cofactors are important for biofilm formation (64, 65). The sulfur relay pathway is involved in the tRNA modifications (66), which are implicated in fitness of bacteria within a biofilm (62). Notably, five additional genes that encode tRNA modification enzymes (*mnmE*, *tilS*, *SUA5*, *dusA*, *tadA*) are inferred to co-evolve with GTA genes (**Table 1**). Finally, the *tadB* gene is involved in the assembly of the tad pili, which have been shown to be involved in promoting the surface colonization and biofilm cohesion (67, 68). Interestingly, in *D. shibae tad* genes are co-regulated with GTA genes, further corroborating linkage between tad pili and GTAs (33).

## Discussion

By showing that GTA head-tail cluster genes, and especially genes within the same functional module of the cluster, tend to co-evolve with each other, and by examining function-coevolution relationship among proteins encoded in a model marine bacterium, we demonstrated that the ERC method is an effective approach to uncover functional relationships among protein-coding genes in bacteria, extending the method’s applicability beyond eukaryotes. Applying the method to GTA-encoding alphaproteobacterial genomes that span >700 million years of evolution (13), we detected a significant evolutionary rate covariation of GTA head-tail cluster genes with 59 protein-coding genes. Multiple genes in this dataset are involved in stress response, DNA repair, homologous recombination, and biofilm formation. These functions are consistent with the accumulating experimental and computational evidence about GTA production, regulation and function in *R. capsulatus* and *C. crescentus*, and with previous hypotheses and models of GTA production triggered under environmental stress (6, 20), GTAs being involved in DNA repair in recipient cell (6, 69), and, most recently, of GTA production occurring in biofilms (36). Our discovery that 13 biofilm-implicated genes co-evolve with GTA genes further highlight the potential importance of biofilm settings for GTA production.

Alphaproteobacteria in general, and GTA-producing *R. capsulatus* and *C. crescentus* in particular, are known to form biofilms (36, 70–72). Biofilms provide a microbial community with benefits that cannot be achieved by the individual cells, such as protection against antibiotics (73) and viral infections (74, 75). Despite either shown or hypothesized benefits of GTA-disseminated DNA to the recipient cells (2, 6, 76), zero relative fitness of the lysed GTA-producing cells implies that the trait of encoding GTAs can only be favored by selection when the benefits of receiving GTAs are confined to either clonal or very closely related cells that also possess the genes encoding the suicidal GTA production trait. Encoding genes for synthesis of specific polysaccharide receptors, which facilitate the adsorption of GTAs (22) and co-regulating the receptor production in one fraction of a population with GTA production in another fraction of a population, can limit GTAs’ targets only to clonal cells or very closely related species. Additionally, success of homologous recombination declines exponentially with the increase in genomic sequence divergence (77, 78), further restricting the usefulness of GTA particles for DNA repair to closely related cells.

Within such groups of closely related cells, GTAs can be viewed as “population-level goods”. However, any population-level goods system faces an inevitable rise of cheaters (79), and a population of GTA producers would be susceptible to cheaters that would not produce GTA particles but still have surface polysaccharides that serve as GTA receptors. Consistent with these conjectures, we observe both the pseudogenization and complete loss of GTA systems in multiple alphaproteobacterial lineages (11, 13, 19). Pseudogenized GTA gene clusters may represent recently emerged cheater lineages, while the absence of GTA gene clusters could be a result of cheater takeover in a species and consequent loss of GTA genes due to the deletion bias (80). (It should be noted that, in both cases, the loss of GTA production in these lineages could also be attributed to acquisition of alternative molecular mechanisms to cope with nutritional stress and DNA repair, or inhabiting niches where GTA production costs outweigh its benefits.) Mathematical modeling showed that maintenance of GTA production trait can be difficult in mixed populations of GTA producers and cheaters, at least under some conditions (81). Yet, despite the likely appearance of cheaters and an observation of recurrent GTA loss in multiple lineages, the GTA production trait has been persisting in alphaproteobacteria for hundreds of millions of years (13), suggesting that some kind of population-level selection is successful. However, details of how such selection operates remain unknown.

One possible solution for an “altruistic” trait to persist in a population over time is to have the population segregated into small sub-populations, an evolutionary scenario first modeled by Wilson (82) and subsequently shown to be equivalent to multi-level selection models that emphasize close relatedness of sub-population members (83, 84). Under this model, the cheaters arise stochastically and therefore are found in many but not all sub-populations. Notably, sub-populations without cheaters outcompete sub-populations with cheaters by having overall higher total productivity due to benefits of the population-level goods produced by the altruistic trait. We hypothesize that GTA systems persist over time because GTA production occurs in such spatially structured populations. We further hypothesize that *it is biofilms* that facilitate the division of GTA-producing populations into isolated sub-populations. A biofilm would ensure that clonal or closely related cells are in spatial proximity, plus they would protect the GTA-producing sub-population from being invaded by other cells, including cheaters. Dense packing of cells would also ensure that the released GTA particles are not dispersed in the environment and reach their recipients (36). Biofilms would also trap the organic debris of the lysed GTA-producer cells within the biofilm (85). DNA of these lysed cells could be used as a part of a biofilm matrix (85), enhancing the sub-population isolation and protection from environmental hazards (86). Other environmental and cellular debris could increase localized nutrient availability. Previous experimental work on biofilms supports some of our conjectures. In the GTA-producing *Caulobacter crescentus* the biofilm structure promotes clonal cells to reside in proximity (71). Biofilm formation is also increased during the ecological competition, providing microbes with the protection to resist invasion by different strains (87).

Building on the current knowledge and previous models of GTA evolution and function, we propose the following model of selection acting on a structured, biofilm-forming bacterial population and maintaining GTA production (**Figure 5**). When a population experiences starvation, nutritional stress increases the generation of reactive oxygen species, which induces DNA damage (88). In a structured population, the fate of each sub-population (i.e., of individual biofilms) depends on the genetic make-up of its cells. In a sub-population of GTA producers (GTA^+^ cells), a small fraction of cells is “sacrificed”, and their DNA is delivered to the remaining cells or hoarded as part of the biofilm matrix, while the cellular debris are utilized as additional nutrients. The resources are protected from invaders by the biofilm structure. As a result, the sub-population counteracts the negative effects of nutrient scarcity and DNA damage, and thus experiences, at worst, only a small population decline due to the lysed GTA^+^ cells. However, if cheaters arise within the biofilm boundaries, the composition of such sub-population under multiple episodes of nutritional stress will change to a higher fraction of GTA non-producers (GTA^-^ cells), as these types of cells will not be lost due to lysis during GTA production and yet will experience all benefits of the population-level goods released by GTA^+^ cells. Eventually, the sub-population will lose GTA production trait. In a sub-population of GTA^-^ cells, mutational load due to DNA damage and limited nutrients will result in a decline of the sub-population size. Therefore, GTA^+^ sub-populations will have higher relative fitness in comparison to GTA^-^ sub-populations, resulting in the larger overall number of GTA^+^ cells in the combined population. Thus, the GTA production trait will be maintained in the population as a whole.

**Figure 5.**
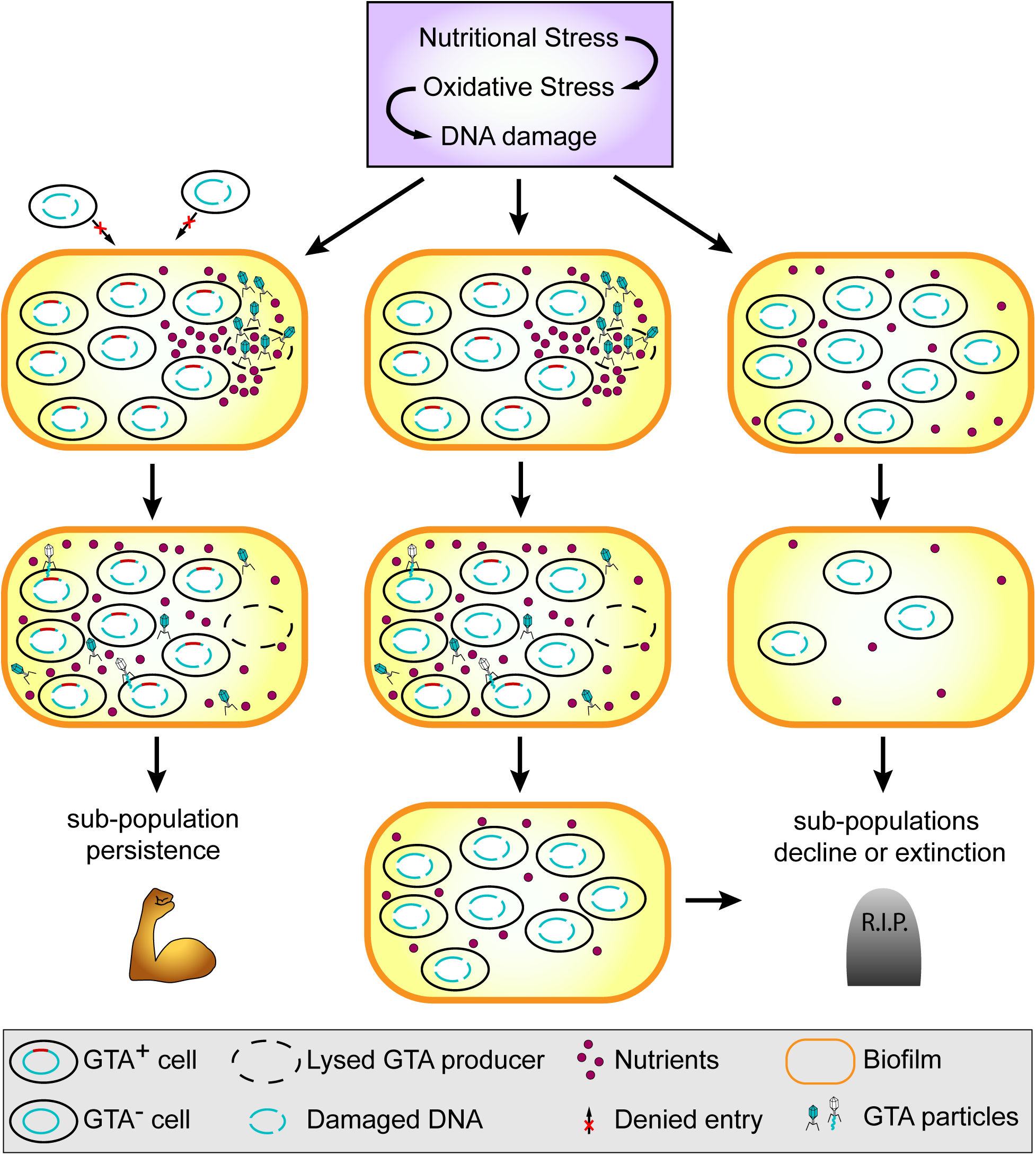
The proposed model of between-group selection that preserves the trait of GTA production in a bacterial population of closely related cells. See **Discussion** section for detailed description.

Although bits of experimental evidence used in the above model come mostly from the research on GTAs in *Rhodobacter capsulatus* and *Caulobacter crescentus*, we hypothesize that our model is applicable to other GTA-containing alphaproteobacterial species, because the detected co-evolution patterns span multiple diverse alphaproteobacterial clades.

## Materials and Methods

### Identification of 293 representative alphaproteobacterial genomes with GTA regions

Initially, 1,642 alphaproteobacterial genomes and annotations of their protein-coding genes were retrieved from the NCBI’s Assembly and RefSeq database (accessed June 2022) (89) (**Table S2**). In these genomes, GTA regions were predicted using GTA-Hunter program with default parameters (19). Because GTA-Hunter looks only for 11 out of the 17 genes of the RcGTA’s head-tail cluster, the remaining GTA genes were identified via BLASTP searches (E-value < 0.1) (90), using as queries the curated set of GTA regions from (19). Only BLASTP matches that are located within the GTA-Hunter-predicted GTA regions were added. Using this procedure, GTA regions were identified in 701 genomes.

To avoid presence of multiple highly similar GTA regions in downstream analyses, the 701 genomes were clustered into 392 Operational Taxonomic Units (OTUs) using the <u>A</u>verage <u>N</u>ucleotide <u>I</u>dentity (ANI) cutoff of 95%, calculated via fastANI v1.1 (91). Within each OTU, GTA regions were examined for “completeness”, defined as having 14 out of the 17 head-tail cluster genes (genes *g1*, *g3.5* and *g7* were excluded because they are not easily detected across GTA-contaning alphaproteobacterial clades (13)). Incomplete GTA regions were discarded. This criterion reduced the number of OTUs to 293. Within each of the 293 OTUs, only one, randomly selected, genome and its GTA region were retained for subsequent analyses (**Table S3**).

### Reconstruction of the reference phylogenomic tree

From the set of 120 marker genes widely used for phylogenomic taxonomy (92), 84 gene families were detected in a single copy in at least 95% of the 293 genomes using AMPHORA2 (93). Amino acid sequences of each of these 84 gene families were aligned using MAFFT v7.505 with the ‘linsi’ option (94). The alignments were concatenated, and the best substitution model for each alignment and the optimal partition scheme were established via ModelFinder (95). The maximum-likelihood phylogeny was reconstructed using IQ-TREE v2.2 (96). The tree was rooted using the *Emcibacterales* and *Sphingomonadales* taxonomic orders, using the previously observed branching order of the *Alphaproteobacteria* as a guide (13, 19).

### Selection of Reference GTA genes

The amino acid sequences of the 14 GTA genes from the GTA regions of the 293 genomes were retrieved and aligned using MAFFT v7.505 with the ‘linsi’ option. Phylogenetic trees were reconstructed from each alignment using IQ-TREE v2.2 (96) under the best substitution model identified by ModelFinder (95). Each tree was compared to the reference tree using the normalized quartet scores calculated in ASTRAL v5.7.8 (97). Eleven GTA genes that exhibited a high congruency with the reference phylogeny (quartet score > 0.8) (**Figure S2**) were designated as “reference GTA genes” and are referred as such throughout the manuscript.

### Identification and functional annotation of gene families in 293 GTA-containing genomes

Gene families were defined as orthologous groups identified in Broccoli v1.2 (98), using DIAMOND (99) for protein similarity searches and the maximum-likelihood method for phylogenetic reconstructions. Gene families that are present in a single copy in at least 50% of the 293 GTA-containing genomes (1,470 in total) were retained for further analyses.

To assign COG functional annotations to the gene families, one randomly selected representative from each family was used as a query against the eggNOG database v5.0.2 and processed through eggNOG-mapper v2.1.9 workflow (100). For gene families found to be co-evolving with the GTA region (see below), additional annotations were sought out using PaperBLAST (accessed in December 2022) (101) and CD-searches against CDD database v3.20 (accessed in December 2022) (102).

### Inference of evolutionary rate covariation

Amino acid sequences of each gene family and each reference GTA gene were aligned using MAFFT v.7.505 with the ‘linsi’ option (94). For gene families that are not found in all 293 genomes, the absent taxa were pruned from the reference tree using functions from the *ete3* package (103). For each gene set, the topology of taxa relationships was constrained to the reference phylogeny and branch lengths were estimated via IQ-TREE v2.2 (96), using the best substitution model suggested by ModelFinder (95). The trees were rooted using relationships in the reference phylogeny as a guide.

Covariations of evolutionary rates among 1,470 gene families and 11 GTA reference genes were examined rates using the CovER pipeline, as implemented in PhyKIT v1.11.12 (38, 104). Within the pipeline, the following steps were carried out. For each pair, their trees were pruned to retain only shared taxa. All trees were corrected for the differences in mutation rates and divergence times among taxa; this was accomplished by dividing the length of each branch by the length of the corresponding branch of the reference tree. Branches with the normalized length > 5 were removed from further analyses, and the retained branch lengths were Z-transformed. For every pair, Pearson correlation coefficient was calculated. A pair of genes was designated as co-evolving, if the Pearson correlation coefficient was positive and had p-value < 0.05 after Bonferroni correction for multiple testing.

The above-described co-variation analysis was carried out on five datasets: among 11 GTA reference genes (11 x 10/2 = 55 comparisons), between GTA genes and 1,470 gene families (11 x 1,470 = 16,170 comparisons), among 1,320 gene families present in GTA-containing genome *Phaeobacter inhibens* (1,320 x 1,319/2 = 870,540 comparisons), among 1,199 gene families present in GTA-producing genome *Caulobacter crescentus* (1,199 x 1,198/2 = 718,201 comparisons), and among 1,255 detected in GTA-producing genome *Dinoroseobacter shibae* (1,255 x 1,254 = 786,885). The *P. inhibens, C. crescentus and D.shibae* analyses resulted in 10,514, 9,373 and 9,675 co-evolving gene pairs, respectively (**Figure S1**). The gene pairs information was assembled into co-evolution networks, in which nodes represent gene families and edges depict co-evolution relationships. To annotate nodes in the co-evolution networks with functional categories, the sub-networks of 1,040 out of 1,320 gene families (for *P. inhibens*), 953 out of 1,199 gene families (for *C. crescentus*), and 997 out of 1,255 gene families (for *D. shibae*), that had unambiguous COG assignments were extracted. The COG functional annotations of genes were assigned as labels.

To minimize the number of false positives and to shorten the list of candidate gene families that co-evolve with GTA genes, the following criteria were applied in addition to Pearson’s R > 0 and Bonferroni-corrected p-value < 0.05: A gene family was required (i) to co-evolve with at least 5 GTA reference genes and (ii) to be above the 95th percentile in the of 1,470 gene families ranked by both p-value and Pearson’s R for at least 5 GTA reference genes. Under these criteria, 59 gene families were retained for further analyses.

### Reconstruction of protein-protein interaction networks

Amino acid sequences of the 1,320 (*P. inhibens*), 1,199 (*C. crescentus*), and 1,255 (*D. shibae*) above-described gene families were used as queries against the STRING database v.11.5 (45), with the high confidence score cutoff and all available sources. This search resulted in 8,612, 8,385, and 8,366 interacting protein-protein pairs for *P. inhibens*, *C. crescentus*, and *D. shibae*, respectively. The information was assembled into a *Phaeobacter inhibens* protein-protein interactions (PPI) networks, in which nodes represent genes and edges depict PPIs.

### Reconstruction of co-fitness networks

*P. inhibens*’, *C. crescentus’*, and *D. shibae*’s gene pairs with similar fitnesses across a wide range of different experimental conditions were retrieved from the Fitness Browser (46) (accessed October 2023). Gene pairs were designated as co-fit if they either had a co-fitness value > 0.75, or they had a co-fitness value > 0.60 and were conserved in other bacterial species (46). This search resulted in 569, 2,657, and 540 co-fit gene pairs for *P. inhibens*, *C. crescentus*, and *D. shibae*, respectively. The information was assembled into co-fitness networks, in which nodes represent genes and edges depict co-fitness associations.

### Comparison of networks and identification of subnetworks

The similarity between *P. inhibens’ C. crescentus’*, and *D. shibae*’ co-evolution networks and either PPI or co-fitness networks was assessed by calculating Jaccard index (105), which measures the fraction of edges shared between the networks. The null distribution of Jaccard indices was created by randomly re-shuffling of the evolutionary rate covariation network 1,000 times.

Tendency of nodes from the same COG category to connect with each other was measured by an assortativity coefficient calculated for the co-evolution subnetworks with unambiguous COG assignments. The coefficient quantifies the connectivity of nodes belonging to the same class and varies between 1 (only nodes of the same class are connected) and –1 (only nodes that belong to different classes are connected). To assess the significance of the observed assortativity values, the permutation test was performed by random shuffling of COG labels 1,000 times. The values of assortativity coefficients were calculated using *igraph* v1.3.5 (106).

All networks were visualized using *igraph* v1.3.5 (106).

### KEGG pathways enrichment analysis for 59 genes that co-evolve with GTAs

Each of the 59 gene families was assigned a KEGG Orhology (KO) label using BlastKOALA (107). Significantly enriched pathways were identified by the hypergeometric test (p-value < 0.05 with Bonferroni correction for multiple testing), as implemented in the *clusterProfiler* package v4.4.4 (108).

## Data Availability

The genomes used in this study are publicly available via NCBI Assembly (https://www.ncbi.nlm.nih.gov/assembly) database. The accession numbers of these genomes are listed in **Table S2**. The following datasets, which were derived from the genomes, are available in the FigShare repository (https://doi.org/10.6084/m9.figshare.23929551.v1): multiple sequence alignment of GTA genes; unconstrained and constrained phylogenetic trees of GTA genes; concatenated alignment of phylogenomic markers and reconstructed reference phylogenomic tree; amino acid sequences of 1,470 gene families; constrained phylogenetic trees of 1,470 gene families; covariation of evolutionary rates for all performed pairwise gene comparisons; *Phaeobacter inhibens*’, *Caulobacter crescentus’, Dinoroseobacter shibae’*s co-evolutionary networks and their subnetworks of nodes with unambiguous GOG assignment, protein-protein interaction networks, fitness networks, and lists of COG functional category assignments for nodes.

## Acknowledgements

We thank Dr. Carey D. Nadell (Dartmouth College) for stimulating discussions about biofilms and social evolution theory, and for critical reading of the manuscript. The work was supported in part by Dartmouth Fellowship and Cramer funds to RK, and Dartmouth Dean of Faculty funds to OZ.

