## Supplementary Figures S1, S2 for "Co-evolution of gene transfer agents and their alphaproteobacterial hosts"

for

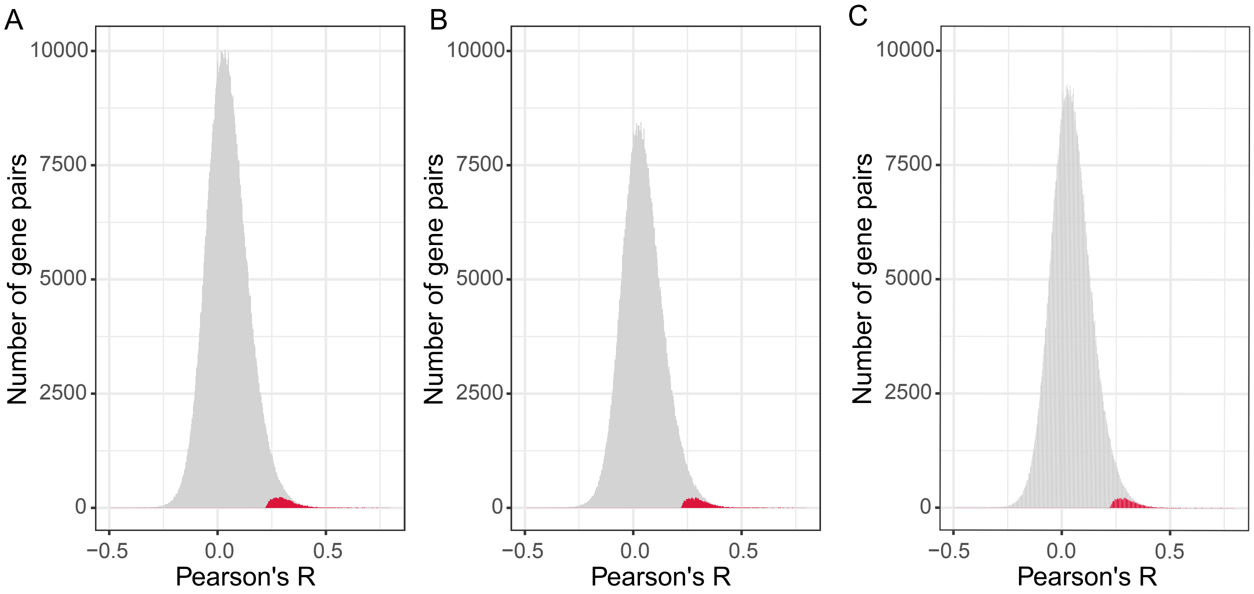


**Figure S1. Pearson's correlation coefficients in pairs of genes from (A) *Phaeobacter inhibens*, (B) *Caulobacter crescentus*, and (C) *Dinoroseobacter shibae* genomes.** Significantly co-evolving gene pairs (Pearson's R > 0 and p-value <0.05 after Bonferroni correction) are highlighted in red.

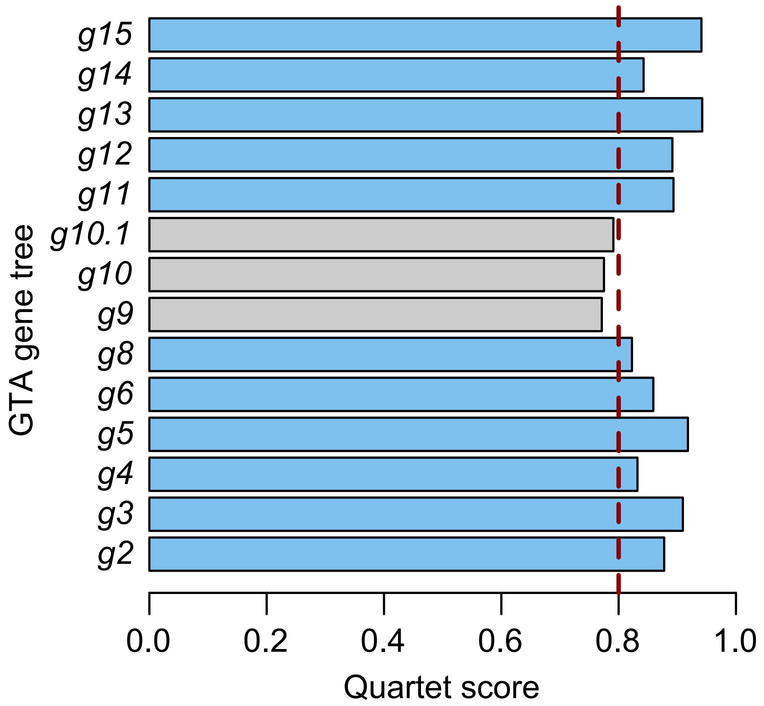


**Figure S2. Comparison of GTA head-tail phylogenies and the reference phylogenetic tree.** The trees were compared using the quarte score metric (X axis). GTA genes are designated using the gene names of the RcGTA. Dashed red line represent the score cutoff used to define “high congruency”. Bars in blue color highlight phylogenies of the reference GTA genes.
